# Role of Immune Checkpoint Proteins in Idiopathic Pulmonary Fibrosis

**DOI:** 10.1101/173237

**Authors:** David M Habiel, Milena Espindola, Chris Kitson, Anthony Azzara, Ana Lucia Coelho, Cory M Hogaboam

**Author notes:** Correspondence, Cedars-Sinai Medical Center, 127 S San Vicente Blvd., AHSP A9315, Los Angeles, CA 90048 [**Author Approval**: All authors have seen and approved this manuscript.].

## Abstract

Idiopathic pulmonary fibrosis (IPF) is a fibrotic lung disease, with unknown etiopathogenesis and suboptimal therapeutic options. Due to the lack of clinical efficacy of standard immuno-suppressants in IPF, the role of the immune response in this disease remains elusive. Nevertheless, previous reports have shown that increased T cell numbers and phenotype is predictive of prognosis in IPF, suggesting that these cells might have a role in this disease. Transcriptomic analysis of CD3^+^ T cells isolated from IPF lungs removed prior to lung transplant (i.e. explant lung) revealed a loss of CD28 expression and both elevated checkpoint and lymphocyte activation pathways. Flow cytometric analysis of a mixture of immune and non-immune cells isolated from explanted IPF lungs showed elevated PD-1 and CTLA4 protein expression on CD4^-^ lymphocytes and PD-L1 expression on EpCAM^+^ and CD45^-^ EpCAM^-^ cells. Lung remodeling and loss of BAL surfactant protein C were observed in NOD SCID IL-2R_*γ*_^-/-^ (NSG) mice that received an intravenous injection of a mixture of IPF cells, including purified IPF T cells. Finally, in humanized NSG mice, anti-CTLA4, but not anti-PD1, mAb treatment induced an expansion of CD3^+^ T cells and accelerated lung fibrosis. Together, these results demonstrate that IPF T cells are profibrotic but the immune checkpoint protein, CTLA-4, appears to limit this effect in IPF.

## Introduction

Despite the advent of approved pharmacological interventions, IPF remains one the most challenging interstitial lung diseases to manage clinically^1^. This lung-localized disease is characterized histologically by the presence of usual interstitial pneumonia, and specifically by the presence of fibroblastic foci, which are believed to be the site of active tissue remodeling. The fibrotic triggers in IPF are unknown but it is speculated that persistent lung injury leads to alveolar epithelial cell injury and death, and subsequent aberrant repair mechanism(s) ablates the alveolus^2^. The nature of the factors dictating these aberrant repair mechanisms in IPF are unknown.

Experimental evidence and histological analysis indicate that there are increased numbers of innate and adaptive immune cells (particularly lymphocytes3-7), which might disrupt normal lung repair in IPF. Peripheral blood IPF T cells exhibit transcript and/or a surface protein expression signature characterized by a loss of one or more costimulatory molecules, including CD28 and ICOS receptors^4,8^. Indeed, the surface expression of the co-stimulatory molecules CD28 and ICOS are lower in IPF relative to normal peripheral blood CD8^+^ cytotoxic but not CD4^+^ helper T cells^9^. Likewise, the accumulation of CD3^+^ T cells and CD20^+^ B cells in lymphocyte aggregates is well documented in the IPF lungs^7^. Further, there is impairment of Semaphorin 7a-expressing regulatory T cells in bronchoalveolar lavage (BAL) and lung explants from IPF patients^5,6^. Finally, there is a high prevalence of monoclonality and oligoclonality of T cells in IPF, suggesting that these cells might be active in the lungs of these patients^10^-^13^.

In this report, a detailed characterization of the phenotype and function of IPF lung-derived T cells is provided. Compared with normal peripheral blood T cells, IPF T cells from lung tissue express markedly less CD28 while also expressing transcripts encoding for checkpoint pathways and T cell activation. Indeed, immune checkpoint proteins, PD-1 and CTLA-4, are strongly expressed in CD4^-^ IPF lymphocytes relative to their normal counterparts. Further, EpCAM^+^ and CD45^-^ EPCAM^-^ cells express markedly greater amounts of immune checkpoint ligands in IPF relative to normal lungs. The intravenous introduction of IPF cells, including isolated IPF lung T cells, into NSG mice caused lung fibrosis that was evident approximately 65 days after IPF cell administration. Interestingly, normal T cells did not induce lung fibrosis in NSG mice and a notable difference between NSG mice humanized with IPF or normal lung T cells was a loss of BAL surfactant protein C protein in the former group of humanized NSG mice. Finally, anti-CTLA4 mAb, but not anti-PD1 mAb, treatment of humanized NSG mice exacerbated pulmonary fibrosis, which correlated with an expansion of IPF T cells in these mice. Collectively, these results demonstrate that T cells present in IPF lung have the potential to drive fibrotic mechanisms but CTLA-4 expression might impede this process.

## Results

### Enrichment of T cell activation and proinflammatory pathways in IPF lung T cells

Various investigators have reported that T cells from IPF patients are phenotypically and functionally distinct from T cells derived from normal donors^4,8^. Specifically, IPF T cells exhibit monoclonal and oligoclonal properties^10-12^, and react to autoantigens^13^. In the present study, RNAseq data analysis of lung-associated IPF T cells and normal peripheral T cells revealed that there were approximate 1850 transcripts significantly differentially expressed in IPF compared normal T cells **(Figure 1A)**. According to a KEGG pathway analysis, T cell activation pathways were enriched in IPF T cells **(Table S1)**. The top most transcript group enriched in these cells was transcripts encoding for various lysosomal components **(Figure 1B and Table S1)**, which are commonly found in activated T cells^14^. Ingenuity upstream analysis predicted the activation of various pro-inflammatory mediators, (including *TNF*, *IL1*-*β* and *IFN*-*γ*), microbial sensors (including *TLR2*, *TLR3*, *TLR4*, *TLR7* and *TLR9*) and profibrotic mediators (including *PDGF*-*BB*, *TGF*-*β1*, *IL18*, and *IL13)* in IPF relative to normal T cells **(Table S2).** Indeed, there was an enrichment of various transcripts previously shown to be abundantly expressed and/or to play a role in pulmonary remodeling **(Figure 1C).** However, there was an overall reduction in transcripts encoding for the T cell costimulatory protein CD28 **(Figure 1D)**, which was consistent with previous reports^4,8^. Further, there was no evidence for Th2 or Th17 skewing in these cells based upon the lack of transcripts encoding for *IL17*, *IL4*, and *IL13* (data not shown). T cells from one IPF patient (IPF2) showed abundant *foxp3* expression but low *TGF*-*β* expression compared with the other IPF and normal cells analyzed **(Figure 1C-D)**. Thus, these results demonstrate that IPF T cells exhibit an activated genotype.

**Figure 1:**
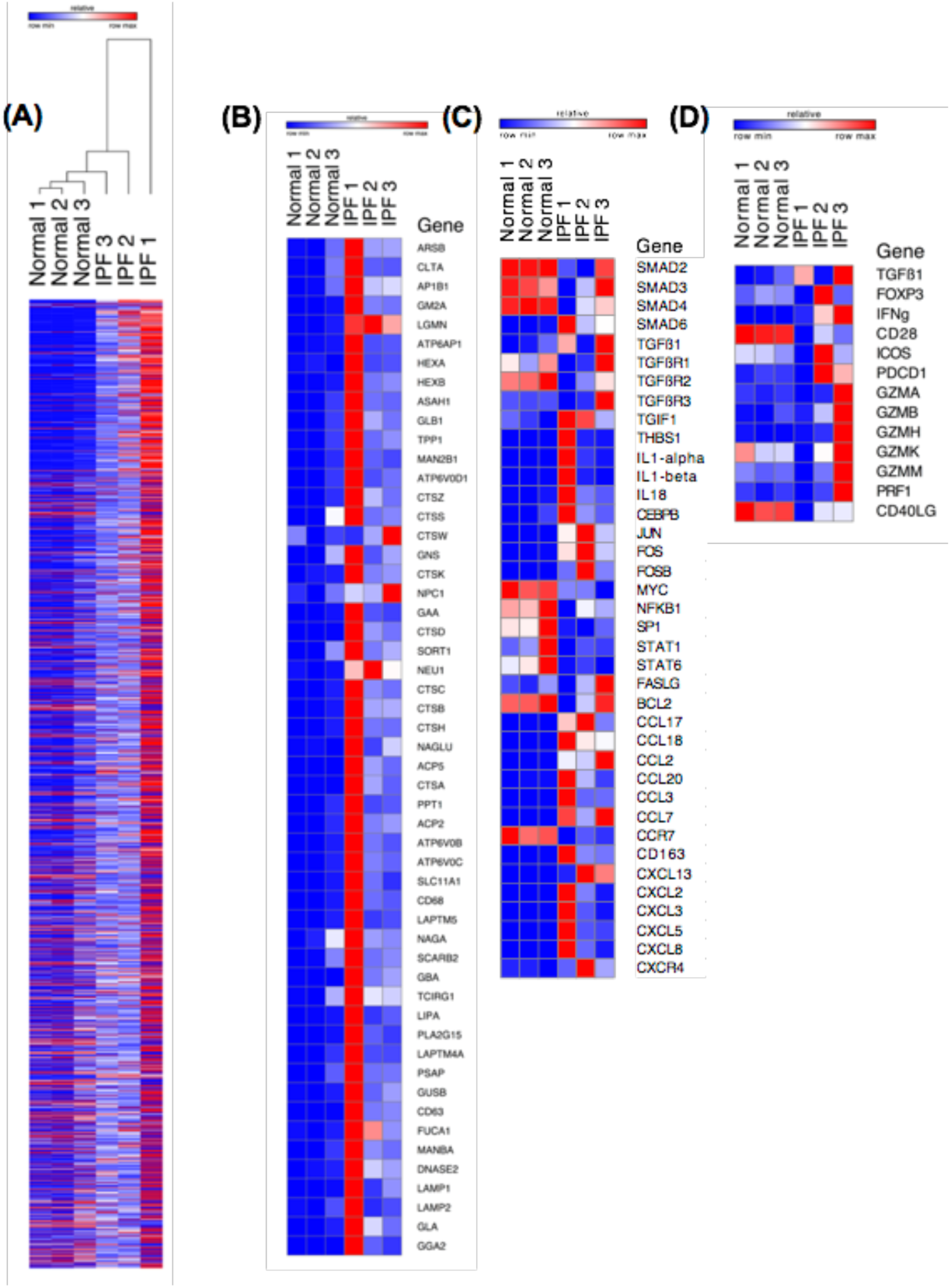
Enrichment of pro-inflammatory and profibrotic transcripts in IPF versus normal T cells. Normal T cells were magnetically sorted from peripheral blood from normal donors and IPF T cells were sorted form IPF lung explant cellular suspensions, RNA was extracted and subject to RNA sequencing. Normalized FPKM values and EDGE significant transcripts were calculated using CLC workbench genomics and imported into GENE-E. **(A)** Depicted are significantly differentially expressed transcripts in IPF relative to normal T cells clustered using one minus Pearson correlation distance metric. **(B-D)** Depicted are heatmaps generated using GENE-E of FPKM values for transcripts encoding proteins involved in lysosomal genesis and function (B) various pro-inflammatory and profibrotic pathways (C) and T cell activation and polarization (D).

### Enrichment of immune checkpoint transcripts in IPF lung T cells

As shown above, IPF T cells appeared to be more active than normal T cells but there was also a loss of co-stimulatory signaling proteins in these cells (^4^,^8,9^, and see above), which is a feature commonly observed in aged and immune checkpoint-regulated T cells^15-18^. Further, there was no conclusive evidence for T cell activation and/or T cell mediated autoimmunity in the IPF patients examined elsewhere or herein. Consequently, we hypothesized that host-mediated immunomodulatory mechanisms such as immune checkpoint pathways were dominant in this disease. Using Ingenuity’s pathway tool, it was noted that there was an enrichment of Immune checkpoint receptors and ligands in bronchial brushings from IPF patients **(Figure 2A)** and isolated IPF T cells **(Figure 2B-C)**. Further, consistent with T cell transcriptomic analysis **(Figure 2B-C)**, CD28 transcript was downregulated in bronchial brushings from IPF versus normal **(Figure 2A)**. Thus, these results indicate that immune checkpoint pathways are elevated in IPF.

**Figure 2:**
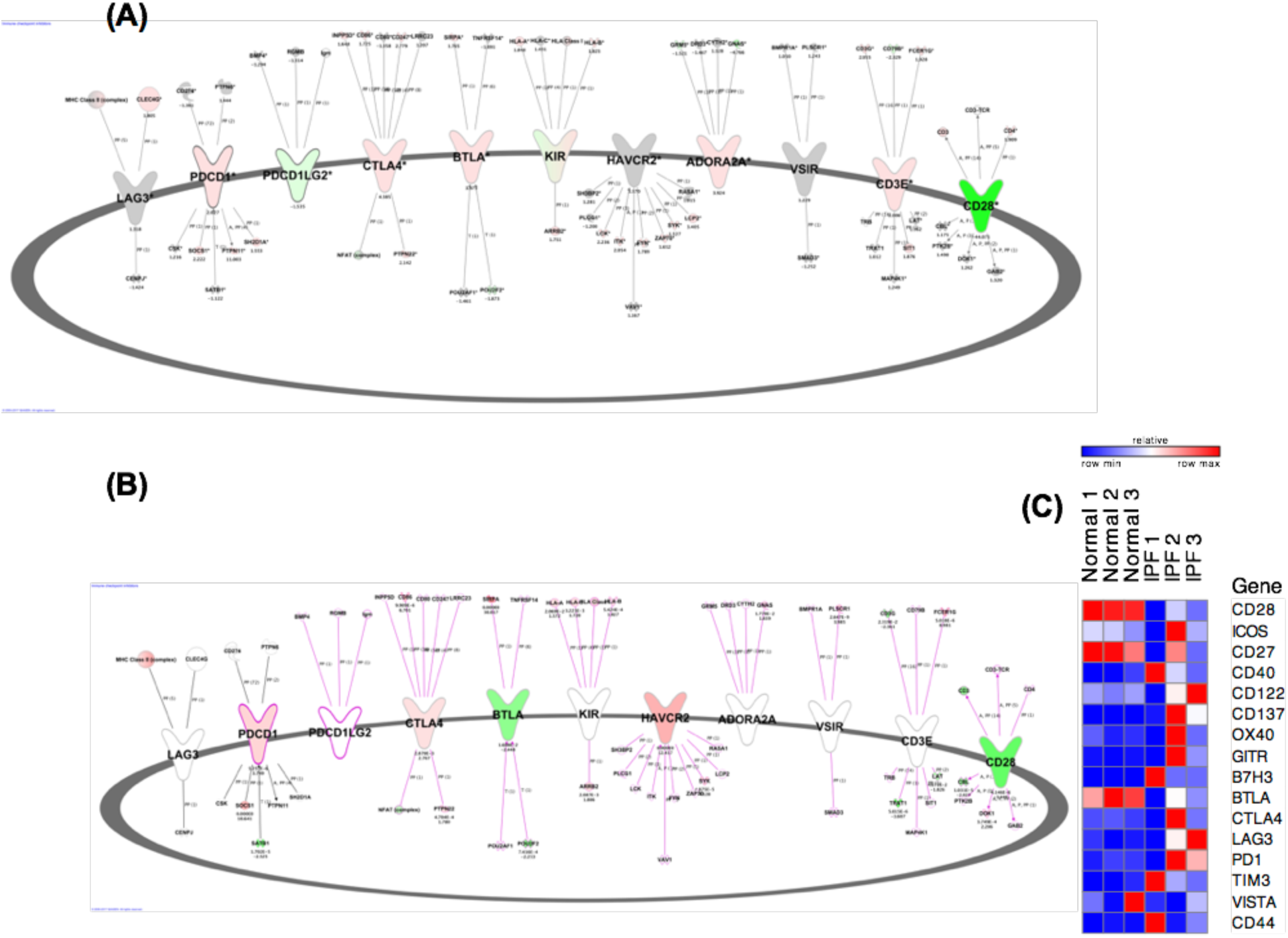
Elevated immune checkpoint protein expression in IPF bronchial brushings and isolated T cells. RNA was extracted from Normal and IPF bronchial brushings and T cells and subject to gene expression array or RNA sequencing analysis, respectively. Ingenuity pathway designer was utilized to generate a custom figure depicting various immune checkpoint, cell surface T cell markers and their directly interacting proteins. **(A-B)** Depicted is an ingenuity custom pathway for immune checkpoint proteins overlaid with gene expression datasets form IPF bronchial brushings (A) or IPF explanted lung derived T cells (B). Transcripts depicted in red and green indicate ≥1.5-fold elevation or reduction, respectively, in IPF relative to normal. Values showing fold change of transcript expression in IPF relative to normal is depicted the top (A) or bottom (B). P-values for expression, whenever available, is depicted on the top (B). Bronchial brushings: n=6 IPF; n=10 normal. T cells: n=3/group. **(C)** Depicted is a heat map generated using GENE-E of FPKM values for transcripts encoding checkpoint proteins in individual normal and IPF samples.

Flow cytometric analysis of immune checkpoint expression in normal, COPD and IPF lung T cells showed that there were no significant differences in the percentage of CD3^+^ **(Figure 3A)**, CD3^+^ CD4^+^ **(Figure 3B)** or CD3^+^ CD8^+^ T cells **(Figure 3C)** between all the patient groups analyzed. Further, there were no significant differences in the percentage of PD-1 or CTLA4 expressing ungated lymphocytes (FSC^low^; **Figure 3D & 3G, respectively),** or FSC^low^ CD4^+^ cells **(Figure 3E & 3H)**. However, there was a significant increase in the expression of both immune checkpoint receptors in FSC^low^ CD4^-^ cells from IPF compared with normal lung samples **(Figure 3F & 3I, respectively)**. Together, these results suggest that there is elevated PD-1 and CTLA-4 protein expression on CD4^-^ lymphocytes in IPF.

**Figure 3:**
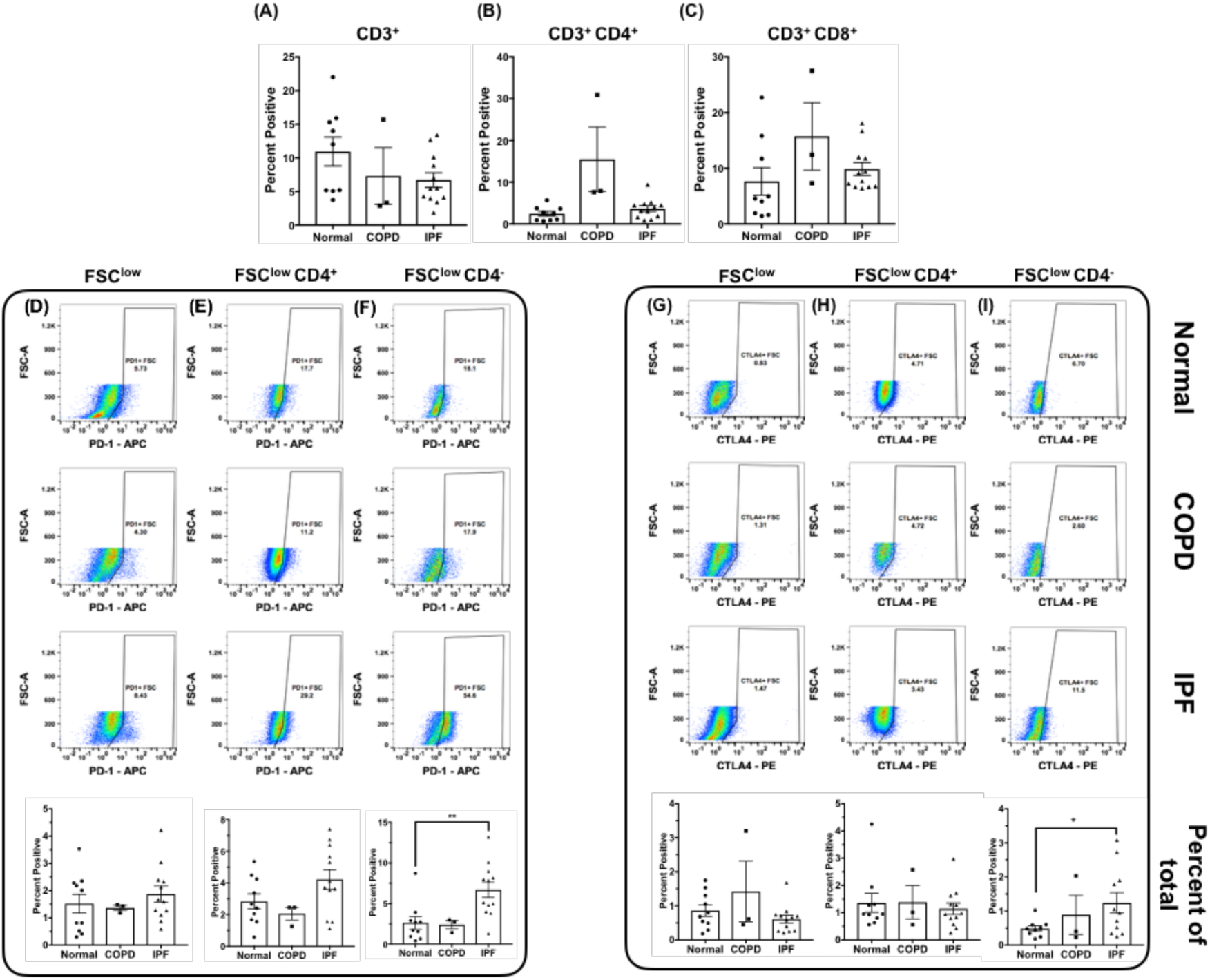
Abundance of PD-1 and CTLA4 expression in IPF CD4^-^ lymphocytes. Flow cytometric analysis of mechanically dissociated normal and IPF lung explant cellular suspensions for cell surface CD3, CD4, CD8, PD-1 and CTLA-4 proteins. **(A-C)** Depicted are bar graphs showing the average percentage of CD3+ (A), CD3+ CD4+ (B) and CD3+ CD8+ (C) T cells in normal, COPD and IPF lung explants. **(D-I)** Depicted are representative dot plots of forward scatter low cells (FSC^low^; D & G), FSC^low^ CD4^+^ (E & H) and FSC^low^ CD4^-^ (F & I) expressing PD-1 (D-F) or CTLA-4 (G-I) from normal (top), COPD (middle) and IPF (bottom) lung explants. Bar graphs (D-I, bottom) depict the average percentage of cells expressing PD-1 (D-F) or CTLA4 (G-I) in normal, COPD or IPF lung explants. n=10-12/group. ^∗^p ≤ 0.05 ^∗∗^p ≤ 0.01

Similar analysis for PD-L1 indicated that this ligand was significantly reduced on CD45+ immune cells in IPF compared with normal lungs **(Figure 4A)**. However, there was a significant elevation in the expression of PD-L1 in CD45^-^ EpCAM^+^ cells from IPF and COPD explants **(Figure 4B)**, and CD45^-^ EpCAM^-^ cells from IPF explants **(Figure 4C)**. Further, there was a significant elevation in the expression of this ligand on CD45^-^EpCAM^+^ cells in IPF compared with COPD explants **(Figure 4B)**. Collectively, these results suggested that unlike PD-1, the expression of PD-L1 was more variable in IPF and appeared to be localized to specific cell types in the IPF lung.

**Figure 4:**
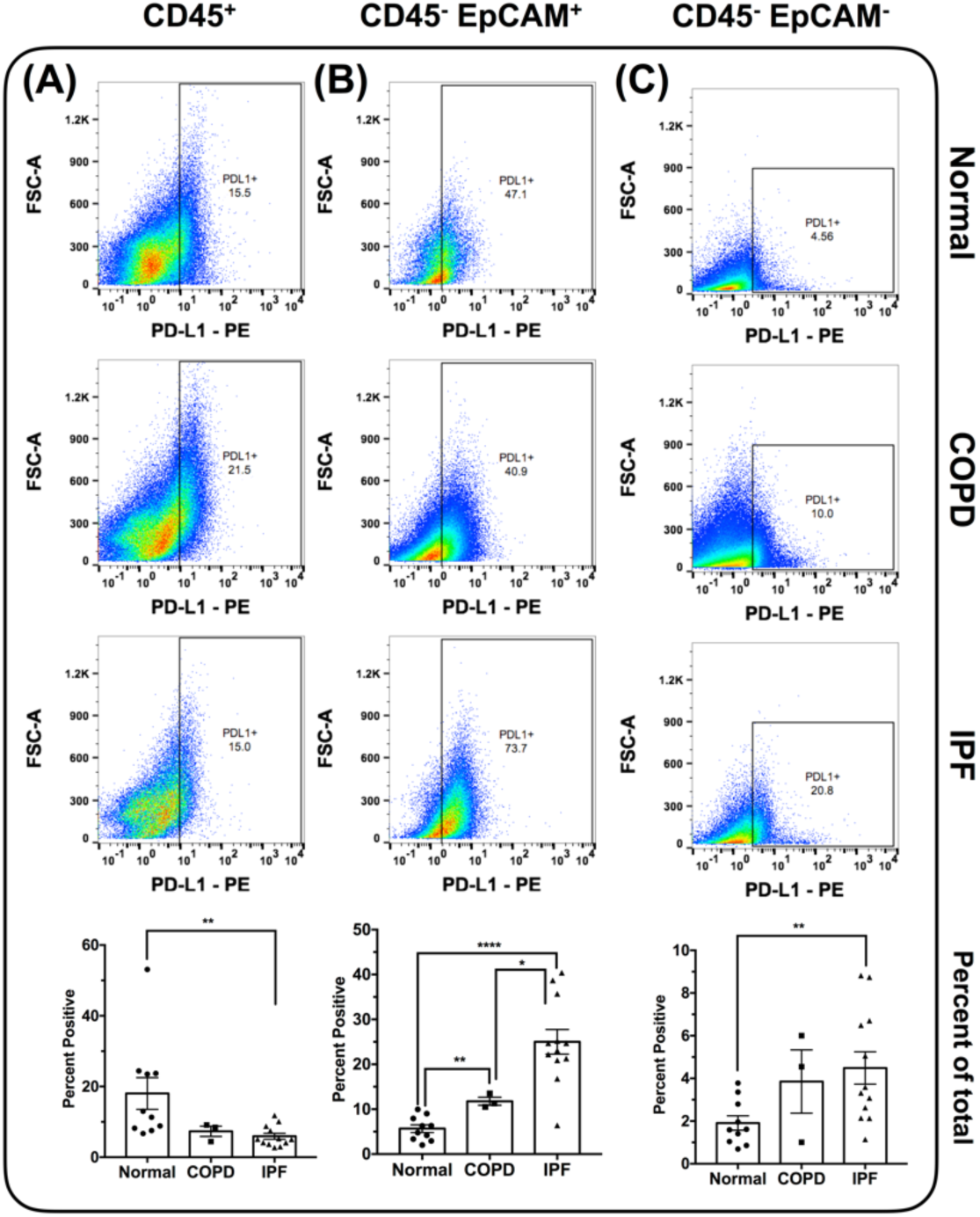
Elevated PD-L1 expression on EpCAM^+^ and CD45^-^ EpCAM^-^ cells in IPF. Flow cytometric analysis of mechanically dissociated normal and IPF lung explant cellular suspensions for cell surface PD-L1 protein. **(A-C)** Depicted are representative dot plots of CD45^+^ (A), CD45^-^ EpCAM^+^ (B) and CD45^-^ EpCAM^-^ (C) cells, expressing PD-L1 from normal (top), COPD (middle) and IPF (bottom) lung explants. Bar graphs (bottom) depict the average percentage of cells expressing PD-L1 in normal, COPD or IPF lung explants. n=10-12/group. ^∗^p ≤ 0.05 ^∗∗^p ≤ 0.01 ^∗∗∗∗^p ≤ 0.0001

### Isolated IPF T cells induced pulmonary fibrosis in humanized NSG mice

Sixty-three days after the intravenous injection of immune and non-immune IPF cells into NSG mice, flow cytometric analysis revealed that human CD3^+^ cells were present in the lungs of these injected NSG mice **(Figure 5A-B and quantified in 5C)** but these cells were more numerous in the spleens of these mice **(Figure 5D-E and quantified in 5F).** Masson’s trichrome staining showed interstitial areas of consolidation and collagenous staining **(Figure 5G-H, blue staining)** in NSG mice humanized with IPF cells. Further, there was a significant elevation in hydroxyproline concentration in the humanized NSG lungs of IPF compared with naïve NSG mice **(Figure 5I)**. Collectively, these studies demonstrate that human T cells engraft in NSG mice after the intravenous introduction of IPF cells into NSG mice and this coincides with lung fibrosis in humanized NSG mice.

**Figure 5:**
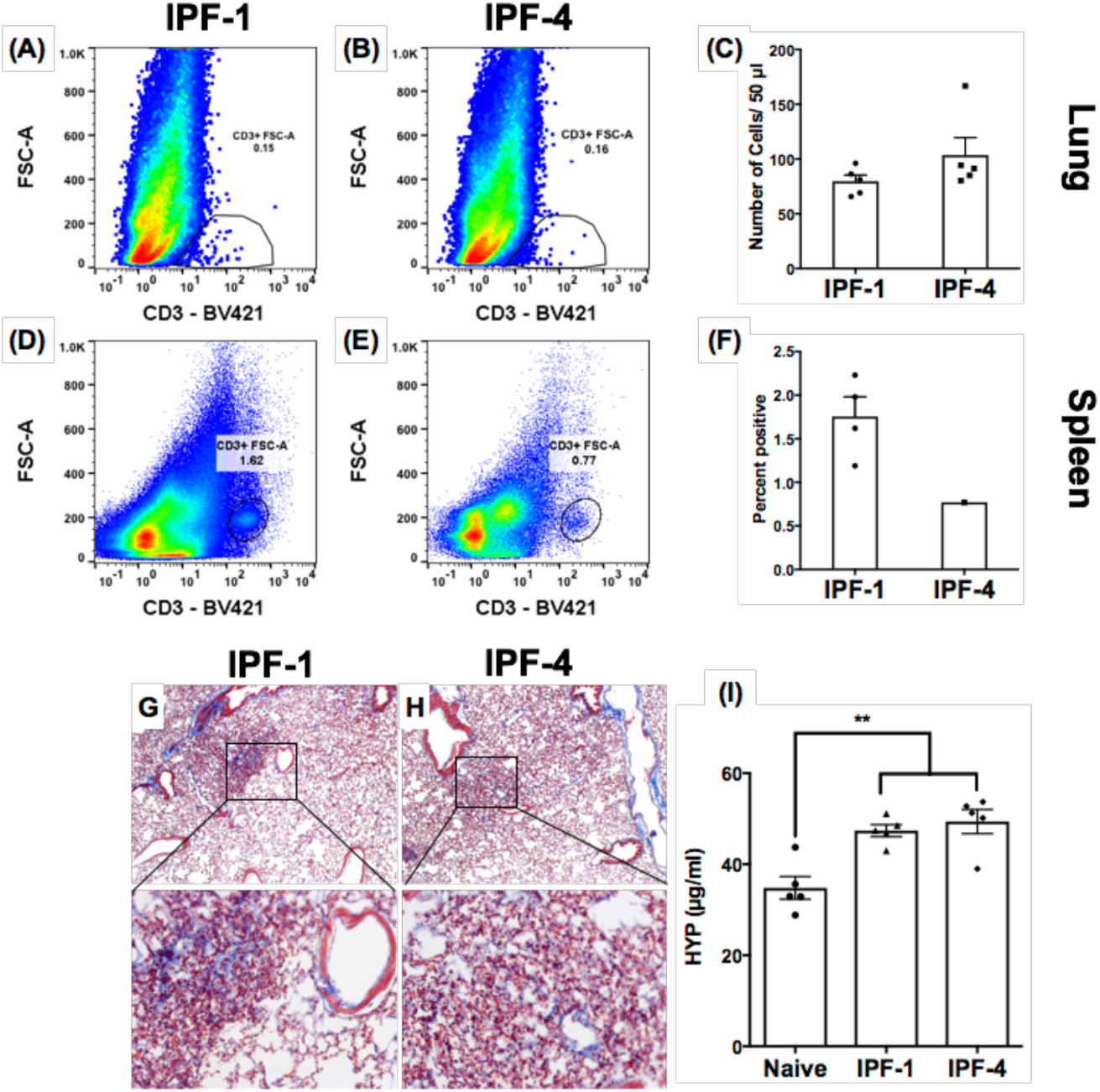
Lung associated IPF T cells engraft in NSG mice. IPF explant cells were IV administered into NSG mice. Sixty-three days after administration, NSG lungs and spleens were collected for flow cytometric, histological and biochemical analysis. **(A-C)** Representative flow cytometric dot plots showing CD3^+^ cells in lungs of NSG mice administered with two different IPF patients (A-B). The average number of CD3^+^ cells in NSG lungs is depicted (C). n=5/group. **(D-F)** Representative flow cytometric dot plots showing CD3^+^ cells in spleens of NSG mice administered with two different IPF patients **(D-E)**. The average percentage of CD3^+^ cells in NSG spleens is depicted (F). n=5/group. **(G-H)** Masson’s Trichrome staining of humanized NSG lungs, 63 days after administration of cells from two IPF patients. **(I)** Shown is the average hydroxyproline concentration in humanized NSG lungs, 63 days post administration of cells from two different patients relative to naïve NSG lungs.

To determine the specific role of T cells in this model, normal and IPF T cells purified from lung samples were intravenously injected into NSG mice. Because of the prognostic importance of CD28 low/null T cells in IPF^4,8,9^, cells from IPF 1 (having the least CD28 and ICOS expression) were used. Sixty-five days after the injection of 1 × 10^4^ T cells from normal or IPF lung explants, Masson’s trichrome staining revealed increased fibrosis in the lungs of NSG mice challenged with IPF T cells **(Figure 6D)** but not in lungs of mice that received normal T cells **(Figure 6B)**, which were more similar to naïve NSG mouse lungs **(Figure 6A).** There was greater picrosirus red staining in the subpleural and interstitial lung regions in NSG mice that received IPF **(Figure 6E)** relative to normal **(Figure 6C)** T cells. Further, there was a significant increase in hydroxyproline in the lungs of NSG mice that receive IPF but not normal T cells **(Figure 6F)**.

**Figure 6:**
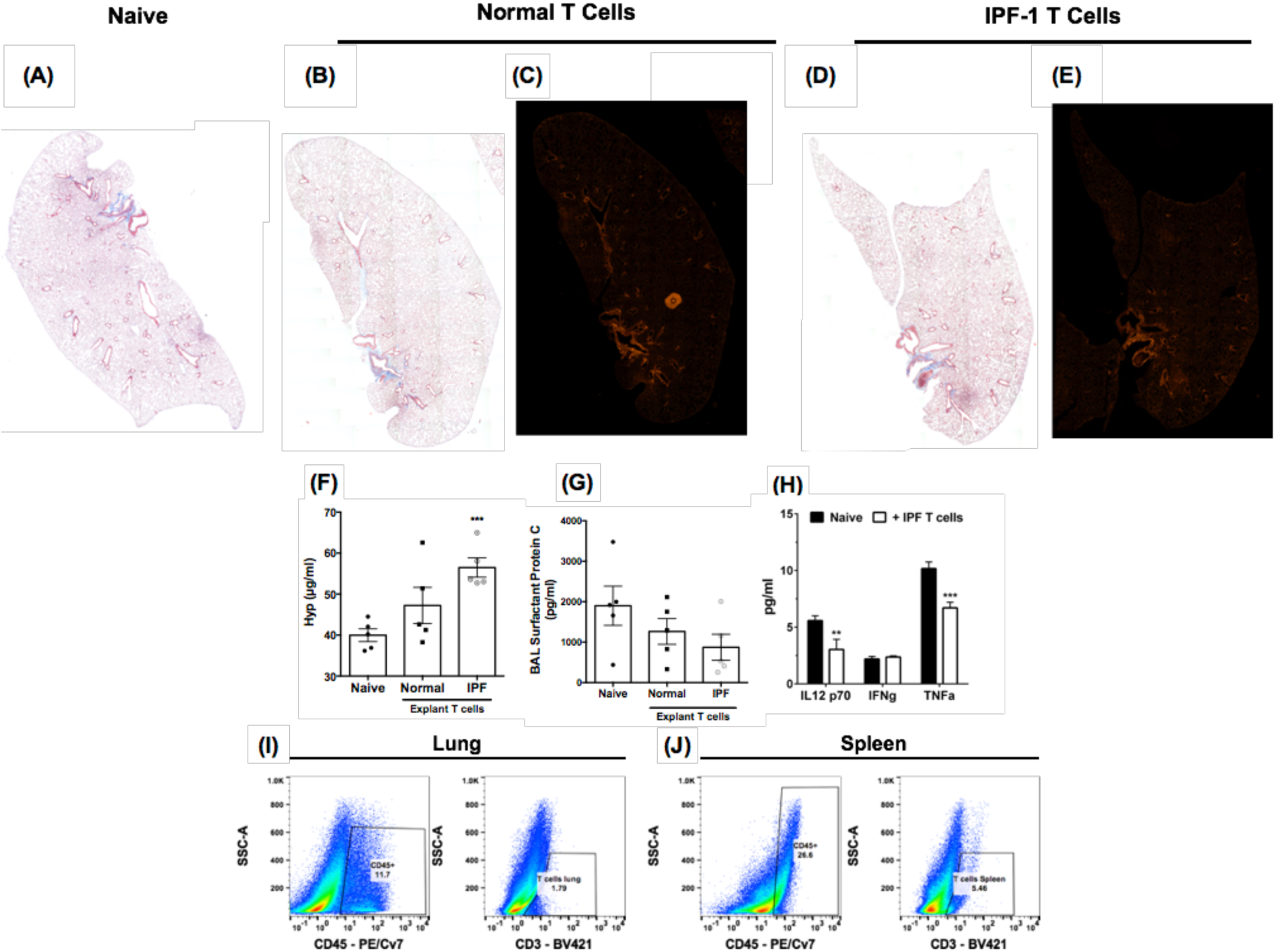
Lung-associated IPF T cells induced lung fibrosis in NSG mice. Normal or IPF lung-associated T cells were FACS or magnetically sorted from lung explants. One hundred thousand cells were intravenously administered into NSG mice and 65 days after injection, mice were sacrificed, their lungs and spleens were collected for histological and flow cytometric analysis. **(A-E)** Depicted are whole mount images of Masson’s Trichrome (A, B & D) or Picrosirius red (C & E) stained lungs from Naïve (A) normal (B-C) and IPF lung-associated T cell (D-E) challenged mice. Shown are representative images from mice challenged with T cells from one normal and two IPF patients. **(F)** Shown is the average hydroxyproline concentration from the lungs of naïve, NSG mice given either normal or IPF T cells. n=5 mice/group. ^∗∗∗^p = 0.0004. **(G)** One milliliter of BAL was collected from naive or T cell challenged mice. Depicted is the average surfactant protein C concentration in the BAL of naïve, normal or IPF T cell challenged NSG mice. **(H)** Depicted is the average IL12-p70, IFN-gamma (IFN*γ*) and TNF-alpha (TNFα) in the BAL from 3-5 mice per group. ^∗∗^p = 0.0063 ^∗∗∗^p = 0.0002. **(I-J)** Shown are representative flow cytometric dot plots depicting human CD45 (left) and CD3 (right) expressing cells in the lungs (left) and the spleens (right) of SCID mice challenged with IPF T cells.

To determine potential mechanisms through which IPF T cells promoted fibrosis in NSG mice, protein analysis of the bronchoalveolar lavage (BAL) fluid was performed for various markers of lung injury and inflammation. Compared with BAL from naïve NSG mice, there was modest decrease in the levels of surfactant protein C in the lungs of NSG mice that received IPF T cells compared with those mice that received normal T cells **(Figure 6G)**. This decline did not appear to be the result of enhanced inflammatory responses because IL12-p70, TNFα and IFN*γ* were modestly reduced or unchanged in the humanized NSG groups compared with the naïve NSG group **(Figure 6H).** Finally, human CD45^+^ cells were detected in the lungs and the spleen of the humanized NSG mice **(Figure 6I-J, left panels, respectively)**, and these cells expressed low levels of surface CD3ε protein **(Figure 6I-J, right panels, respectively)**. Thus, these results demonstrate that IPF T cells appear to be targeting mouse type 2 epithelial cells rather than evoking an inflammatory response in the lungs of humanized NSG.

### Immune checkpoint inhibitors exacerbated pulmonary fibrosis and the expansion of human T cells in humanized NSG mice

We next examined the effects of checkpoint targeting mAbs on the lung fibrosis elicited by the intravenous introduction of IPF cells into NSG mice. Compared with naïve NSG mice **(Figure 7A)**, humanized NSG mice that received anti-CTLA-4 mAb exhibited the greatest histologic evidence of pulmonary fibrosis **(Figure 7D)** compared with saline-treated **(Figure 7B)**, the IgG-treated **(Figure 7C & 7E)**, and the anti-PD-1-treated **(Figure 7F)** NSG mouse groups. Further, while anti-PD-1 mAb did not affect hydroxyproline levels in humanized NSG mice, there was a significant elevation in hydroxyproline in the anti-CTLA-4 mAb-treated group compared with the respective IgG control NSG group **(Figure 7G)**. Anti-CTLA4 mAb treatment increased the numbers of human-CD3^+^ cells **(Figure 7H)** and murine F4/80^+^ CD11c^-^ macrophages **(Figure S2B)** compared with the appropriate IgG control group. Anti-PD-1 mAb treatment in humanized NSG mice increased the numbers CD25^+^ cells **(Figure 7I)** compared with the appropriate IgG control group. Finally, there were no significant changes in the number of human-CD3^+^ **(Figure 7J)** or CD25^+^ **(Figure 7K)** cells in the spleens of any of the NSG mouse groups or in mouse myeloid cells (F4/80^+^ CD11c^+^, F4/80^-^ CD11c^+^ and F4/80^-^ Ly6G^+^ cells) in the lungs of any of the NSG mouse groups **(Figure S2A & S2C-D)**. Thus, these results demonstrate that targeting CTLA4 exacerbated fibrosis in a translational NSG model of pulmonary fibrosis.

**Figure 7:**
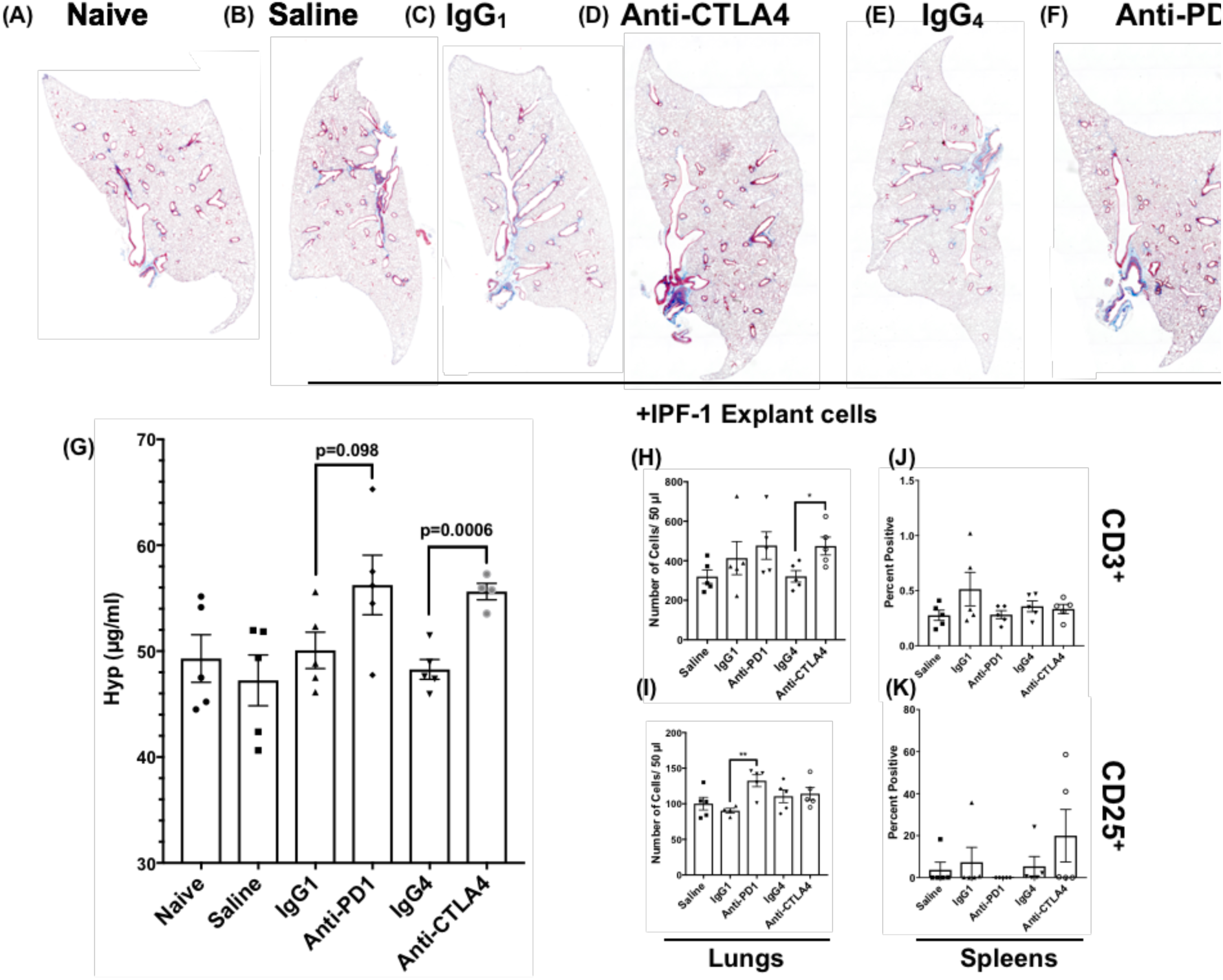
Targeting CTLA-4 in a humanized NSG model of IPF exacerbates lung remodeling. IPF explant cells were intravenously administered into NSG mice. Seven days after administration, mice were treated with anti-PD-1, anti-CTLA4 or IgG control antibodies (5 mg/kg) twice a week for 28 days. After a total of 35 days, mice were sacrificed, their lungs were collected for flow cytometric, histological, and biochemical analysis. **(A-F)** Depicted are whole mount images of Masson’s Trichrome stained lungs from Naïve (A) and IPF cell challenged mice treated with saline (B), IgG_1_ (C), anti-CTLA4 (D), IgG_4_ (E) or anti-PD-1 (F) antibodies. **(G)** Shown is the average hydroxyproline concentration from the lungs of naïve xenografted NSG mice, treated with saline or antibodies. n=5 mice/group. Lung and spleen cellular suspensions were generated from naïve and xenograft antibody treated NSG mice and subject to flow cytometric analysis for human CD45, CCR10, CD3 and CD25. **(H-I)** Depicted is the average number of CD3^+^ (H & J) or CD25^+^ (I & K) cells in the lungs (H-I) and spleens (J-K) of NSG mice. n=4-5/group. ^∗^p ≤ 0.05 ^∗∗^p ≤ 0.01

## Discussion

While the precise role of the immune system on the progression of IPF remains controversial, recent evidence suggest that immune pathways may be aberrantly activated in this disease^19-24^. These findings provided the impetus to further characterize various immune cell populations in IPF lungs and to determine the roles of these cells in disease initiation and progression. The data present herein indicated that T cells isolated from IPF lung samples alone had profibrotic properties *in vivo*, but the profibrotic nature of these cells was largely kept in check by immuno-modulatory molecules like CTLA-4. Targeting CTLA4 correlated with a marked expansion of human T cells and exacerbated pulmonary fibrosis in a humanized mouse model of IPF. The loss of type 2 epithelial cells was one consequence of T cell activation in humanized NSG mice. Thus, our findings presented demonstrate that IPF T cells are profibrotic when released from checkpoint inhibition.

Our transcriptomic analysis of IPF T cells derived from lung samples revealed an enrichment/upregulation of transcripts commonly found in activated T cells, including those involving lysosomal biogenesis and function and active IL4/IL13-driven STAT6 signaling^14,25^. Further, there was a strong induction of transcripts encoding for *JUN*, *FOS*, and *FOSB* of the *AP*-*1* transcription factor complex, which might be a consequence of T cell activation pathways including TCR signaling^26^. Paradoxically, our transcriptomic analysis of T cells revealed a universal down regulation of CD28 transcripts in IPF compared with normal T cells, which is consistent with various published reports documenting the loss of co-stimulatory proteins such as CD28 and ICOS on IPF T cells^4,8,9^. Loss of CD28 on T cells is a feature of antigen experienced, highly differentiated and aged T cells^15,18,27^, a finding that is consistent with the limited IPF T cell receptor repertoire we observed in our transcriptomic analysis (data not shown) and by others^10-13^. Transcriptomic analysis of CD28-low IPF T cells revealed an abundance of T cell activation pathways and an enrichment of T cell mediated KEGG autoimmune pathways. Indeed, several reports have observed an abundance of CD28(null) T cells in inflammatory disorders, including atherosclerosis, chronic viral infection and autoimmunity (reviewed in ^27,28^). This is supported by a study showing the presence of CD28(null) T cells in IPF lungs, which elaborated significantly higher levels of IL1ß, IL-2, IL6, IL8, IFN*γ*, TNF-α, GMCSF, MIP-1ß, IL4 and IL10 proteins relative to their CD28^+^ counterparts after anti CD3 stimulation^8^. Further, the same study observed an associated poor prognosis in IPF patients showing an abundance of CD28(null) T cells. These results suggest that various subpopulations of T cells in IPF may contribute to disease progression.

To determine potential role(s) of IPF T cells in modulating lung remodeling, we used a humanized SCID model that is initiated via the intravenous introduction of a mixture of immune, epithelial, stromal and endothelial cells from IPF lung explants. These cells induced lung remodeling that was studied most extensively at day 63 after cell injection into NSG mice. The majority of human CD3^+^ cells were localized in the spleens rather than the lungs of humanized NSG, possibly reflecting the fact that these immune cells preferentially engrafted a more compliant environment like the spleen or these injected human cells did not need to displace other resident cells in these immunodeficient mice. The profibrotic effects of IPF T cells *in vivo* was observed after approximately 2 months in NSG mice. In these studies, human-CD45^+^ cells (with side scatter properties consistent with lymphocytes) were abundant in the lungs of NSG mice but these cells showed low surface expression of CD3. It is interesting to note that human T cells engrafted better in the lungs of NSG when purified rather than mixed cell preparations were injected. We speculate that these differences may be partially explained by the expression of immune checkpoint proteins by other immune and structural cells in the mixed cell preparations, which may limit T cell proliferation and/or activation in NSG mice.

With the recent success of immune checkpoint inhibitors in treating various solid tumors, immune checkpoint pathways have been an active area of investigation^29-31^. These pathways have been shown to regulate the activation of acquired immune cells, especially T cells via receptor-ligand signaling and soluble mediators^30-32^. The most studied (and clinically targeted) immune checkpoint proteins include PD-1 and its ligands, PD-L1 and PD-L2, and CTLA-4, which are often elevated in chronic infection and cancer (reviewed in ^30-33^). PD-1 is often observed on chronically-activated effector T cells and regulatory T cells whereas PD-L1 is often observed in tumor cells and on immune and structural cells localized in tumor micro-environments^32-34^. PD-1 activation inhibits T cell receptor signaling^33-35^ and effector T cell activation but this receptor promotes the suppressive functions of regulatory T cells^36-38^. CTLA-4 is predominantly expressed by regulatory T cells, where it interacts with CD80 and CD86 thereby inhibiting T cell activation^17,39^. In our studies, both PD-1 and CTLA-4 were expressed on CD4^-^ lymphocytes, and PD-L1 was present on EpCAM^+^ and CD45^-^ EpCAM^-^ cells in IPF and some normal lung samples. When anti-CTLA-4 mAb or anti-PD-1 mAb were used in humanized NSG mice, the former but not the latter mAb significantly increased lung fibrosis compared with the appropriate IgG-treated NSG group. In addition, anti-CTLA-4 mAb treatment was associated with increased human T cell engraftment and/or proliferation compared with the IgG-treated NSG group. Interestingly, the loss of CD28 expression occurs in T cells that strongly express CTLA-4^16^ and lack of changes observed in the anti-PD-1 mAb treated humanized NSG model might reflect the need for CD28 expression by T cells before these cells respond to PD-1 inhibition^40^. The mechanism(s) via which IPF T cells promote lung remodeling requires further investigation. We observed no significant differences in neutrophils or inflammatory mediators in humanized NSG mice compared with naïve NSG mice. There was no evidence of cellular infiltration or injury in the liver or kidney of humanized NSG mice suggesting that the fibrotic response induced by IPF cells in the lungs of NSG mice was not due to graft-versus-host disease. However, lower surfactant C levels were detected in BAL samples from NSG mice that received IPF T cells, which might be an indication of the injurious effect of IPF T cells directed against alveolar type 2 epithelial cells in the NSG mouse lung. Collectively, our results demonstrate that CTLA-4 might protect against fibrosis in IPF via its modulation of T cells poised to initiate profibrotic mechanisms.

Thus, we have characterized IPF lung T cells and explored key functional features of these cells that indicate that while they are profibrotic, their activity might be kept in check by molecules like CTLA-4 in IPF. These results raise the possibility that enhancing the expression or function of CTLA-4 might limit the injurious and/or profibrotic activity of T cells in IPF.

## Materials and Methods

### Isolation of IPF explant cells and T cell sorting

Normal and IPF lung explants were placed into sterile PBS, washed and transferred into fresh PBS. Tissue was minced and spun at 600 × g for 5 minutes. Supernatants were collected with the PBS utilized to wash the explanted lungs (lung wash). The top layer of the pellet enriched in mechanically dissociated cells and red blood cells (RBCs) were strained through a 70-μm strainer, the strainer washed several times with DPBS to separate cells from the minced tissues. This procedure was repeated until the RBCs and dissociated cells were removed from the minced tissue pellet. The dissociated cells were mixed with the lung wash and spun down at 400 × g for 5 minutes. RBCs were lysed using RBC lysis buffer (Biolegend) and cells were then counted and viably frozen down using CryoStor CS10 freezing medium (STEMCELL Technologies Inc.). For *in vivo* experiments, cells were rapidly thawed, washed in serum free medium and intravenously injected into NSG mice as described below. For *in vivo* experiments, cells were thawed out, T cells were magnetically sorted using anti-human CD3 microbeads (Miltenyi Biotech, 130-050-101), cultured on plastic dishes for 2 hours to deplete any contaminating antigen presenting cells, after which 100, 000 cells were intravenously injected into NOD/SCID mice.

### Mice

Six to eight-week old Pathogen free NOD Cg-Prkdc<SCID> IL2rg<Tm1wil>Szi (NSG) were purchased from Jackson laboratories and housed in Cedars-Sinai Medical Center’s high isolation animal rooms. Mice were allowed a minimum of one week to acclimate, and 1 × 10^6^ mixed cells or 1 × 10^5^ purified T cells were then injected intravenously into groups of 5 or more NSG mice. Mice were monitored daily and were sacrificed if there is clear evidence for morbidity (weight loss of more than 20%, loss of fur, paralysis and/or lack of responsiveness when handled). The remainder of the mice were sacrificed after 63-65 days. To determine the role of immune checkpoint pathways in our humanized NSG model, mixed IPF explant cell-xenografted NSG mice were treated with 5 mg/kg of anti-PD-1 (Opdivo), anti-CTLA-4 (Ipilimumab) or IgG control antibodies (IgG_4_ or IgG_1_, respectively) twice a week for four weeks. After a total of five weeks, mice were sacrificed and their lungs and spleens were collected for flow cytometric, histologic and biochemical analysis. BAL fluid was collected for protein analysis, the superior and middle lobes were collected for biochemical hydroxyproline quantification, the inferior lobe and spleens for flow cytometric analysis, post-caval lobe for quantitative PCR analysis and the left lung for histological analysis.

### Detection of IL12-p70, IFN-gamma and TNF-alpha in the BAL

Naïve and T cell challenged NOD/SCID mice were sacrificed and their lungs were lavaged using 1 ml of saline solution. BAL cells were spun down and the supernatants were frozen until analysis. IL12-p70, IFN-gamma and TNF-alpha levels in the BAL was measured using predesigned Bioplex assays (Bio-Rad) as recommended by the manufacturer.

### Histological analysis

Left lung tissue was fixed in 10% NBF solution for 24 hours and subsequently transferred into tissue cassettes and placed into a 70% ethanol solution for a minimum of 24 hours. Tissues where paraffin embedded, sectioned, and stained with Masson’s trichrome. For Picrosirius red staining, slides were deparaffinized and hydrated, stained with hematoxylin for one minute and then for one hour in a Picrosirius red solution (5 grams Sirius red F3B (Sigma-Aldrich) in 500 ml of saturated aqueous Picric acid solution (Sigma-Aldrich)). After staining, slides where washed in two changes of 0.05% Glacial acetic acid solution, dehydrated and mounted. Stained slides where examined using a Zeiss Axio Observer Z1 microscope and the Zeiss Zen 2012 v 1.1.2.0 software (Zeiss).

### Hydroxyproline assay

Total lung hydroxyproline was analyzed as previously described^113^ with a few modifications. Superior and middle lobes were surgically dissected, and placed into 5 ml sterile tubes and flash frozen until all the samples have been collected. On the day of the assay, tissues were thawed at room temperature and 500 μl of distilled water (dH_2_O) was added to the tissues. Tissues were homogenized using a micro-sample homogenizer (Pro Scientific) as follows: the homogenizing tip was washed in 70% ethanol, then dH_2_O and then utilized to homogenize the tissues. The homogenizer was washed in dH_2_O between samples and a new batch of dH_2_O was utilized after 5 samples. After homogenization, samples were transferred into Fisherbrand Borosilicate glass screw capped tubes with a rubber liner (Fisher Scientific) and 560 μl of 12N HCl was added to the homogenized tissues. Samples were capped, vortexed, placed into a preheated oven set to 120 °C and incubated overnight. The next morning, samples were cooled down to room temperature, vortexed and filtered through a 0.45 μm syringe filter. Fifty microliters of the filtered samples were transferred into a 1.5 ml micro-centrifuge tube and evaporated on a heating block set to 100 °C for 2-3 hours. While the samples were incubating, standards to generate a standard curve were prepared by diluting Hydroxyl-L-Proline (Sigma Aldrich) in Acetate-citrate buffer (17g Sodium Hydroxide (Sigma-Aldrich), 36.2 g Sodium Acetate (Sigma-Aldrich), 25g Citric acid (Sigma-Aldrich), 6 ml of Glacial Acetic acid (Thermo-Fisher Scientific) and a final pH of 6 in a total of 500 ml ddH_2_O) into nine different concentrations ranging from 200 μg/ml to 20 μg/ml. After incubation, the desiccated sample pellets were resuspended in 50 μl of Acetate-citrate buffer and all samples and standards (50 μl) were transferred into 5 ml tubes. One ml of Chloramine-T solution (1.128g Chloramine T hydrate (Sigma-Aldrich) dissolved in 8 ml n-Propanol (Sigma-Aldrich) and 8 ml of ddH_2_O and then supplemented with 64 ml Acetate-Citrate buffer) was added to each sample and standard and mixed by gentle vortexing. Samples and standards were incubated at room temperature for 20 minutes and subsequently 1 ml of Ehrlich’s solution (6.75 g 4-(dimethyalamino)-benzaldehyde, 25.95 ml n-Propanol and 13.65 ml of 60% Perchloric acid) was added and all tubes were mixed by vortexing, transferred into a pre-heated oven set at 65 °C and incubated for 20 minutes. Samples and standards were then taken out of the oven and cooled for 10 minutes at room temperature protected from light. 200 μl of samples and standards were transferred into a flat-bottom 96 well plate and absorbance at 550 nm was recorded using a BioTek Synergy H1 microplate reader (BioTek Instruments Inc.). Sample concentrations were calculated using an equation generated from the standard curve.

### RNAseq analysis

CD3^+^ T cells were sorted from IPF explants and CD4+ cells were magnetically sorted from the peripheral blood of normal donors. RNA was extracted from the sorted cells and a minimum of 200 ng of total RNA was used, samples were sequenced using HiSeq 2000 SBS Sequencing v3. Raw reads in FASTQ format were aligned to the UCSC human reference genome (hg38) using CLC genomics workbench V8.5.1, RPKM values were generated and fold change and statistical significance were calculated using the Empirical analysis of DGE (EDGE;^41,42^). The resulting data were analyzed using QIAGEN’s Ingenuity Pathway Analysis (IPA, QIAGEN Redwood City, www.qiagen.com/ingenuity) and DAVID Bioinformatics Resources 6.7^43,44^. Ingenuity IPA was set to only consider changes in gene expression of 1.5-fold or greater and a p value of 0.05 or less. Ingenuity upstream analysis was exported and is provided in Table S3. For KEGG pathway analysis, average transcripts with an RPKM ≥ 100 in IPF T cells were uploaded into DAVID Bioinformatics Resources 6.7 (Huang da et al., 2009a) and KEGG pathway analysis were exported and is provided in Table S1. For transcript expression for various profibrotic, T cell function and DNA repair transcripts, RPKM values from 3 normal and 3 IPF T cell datasets were imported into GENE-E (http://www.broadinstitute.org/cancer/software/GENE-E/), where heat maps depicting the normalized FPKM counts for each transcript were generated and depicted. Finally, T cell-associated markers and immune checkpoint receptors and ligands were compiled using the ingenuity pathway designer tool and transcript expression data from IPF bronchial brushings or T cells (relative to their normal, non-fibrotic, counterparts) were overlaid.

### Analysis of gene expression arrays of IPF and normal bronchial brushings

RNA was extracted from Normal and IPF bronchial brushing. RNA was then hybridized to probes and subject to microarray analysis. The resulting intensities from 6 IPF and 10 normal samples were averaged and the ratios of IPF-to-normal were calculated. The results were then uploaded onto ingenuity IPA for pathway analysis. Ratios where then converted into fold change in Ingenuity IPA and analysis was set to consider transcripts showing a change of 1.5-fold or greater. The resulting analysis was then overlaid onto a custom generated T cell and immune checkpoint pathway interaction network using Ingenuity’s pathway designer.

### Flow cytometry

#### Murine lungs and spleens

Lungs were surgically dissected and placed into 1 ml of sterile complete medium on ice until all the lobes were collected. Murine lung cellular suspensions were generated using mouse lung dissociation kit, C-tubes and a GentleMacs dissociator (Miltenyi Biotech) as recommended by the manufacturer. Spleens were dissociated using a sterile rubber tip of 1 ml syringe plunger and 100 μm 50 ml tube filters (BD bioscience) as follows: isolated spleens were placed in 1 ml of sterile complete medium until all the samples are collected. Sterile 50 ml conical tube filters were placed on top of sterile 50 ml conical tubes and were equilibrated by pipetting 1 ml of calcium and magnesium free DPBS (DBPS; Cellgro) through the filters. Using sterile forceps, spleens were transferred on top of a sterile 50 ml filter. Cellular suspensions were generated by mashing the spleens using a sterile rubber tip of a syringe plunger. The filters containing mashed spleens were flushed with 5 ml of DPBS. Lung and splenic cellular suspensions were spun down and resuspended in 1 ml of RBC lysis buffer (Biolegend), incubated at room temperature for 1 -2 minutes and then equilibrated by adding 25 ml of DPBS to each tube. Cells were spun down and resuspended in flow cytometric wash/staining buffer (DPBS + 2% FBS) in the presence of human and mouse Fc receptor blocking antibodies (Biolegend). Cells were then stained with anti-human-CD3, -CD25 and anti-mouse F4/80, CD11c and Ly6G antibodies (Biolegend) for 20 minutes at 4 °C. Unstained, isotype controls and staining in naïve unchallenged murine lung and spleen suspensions were utilized to gate out any non-specific antibody binding and background fluorescence. Flow cytometric data were acquired within one week of staining using a MACSQuant 10 (Miltenyi Biotech) flow cytometer and data were analyzed using Flowjo software V10.2 (Treestar Inc.).

#### Lung explant suspensions

Explant suspensions were processed as described above, resuspended in flow cytometry wash/staining buffer and blocked with anti-human Fc receptor antibodies (Biolegend) for 15 minutes at 4 °C. After blocking, cells were stained with anti-human CD45, EpCAM, CD3, CD8, CD4, CTLA4, PD-1 and/or PD-L1 antibodies (Biolegend) for 20 minutes at 4 °C. Unstained, isotype and fluorescent minus one controls **(Figure S1A-C)** were utilized to gate out any non-specific antibody binding and background fluorescence. Cells were then washed twice with flow cytometry wash/staining buffer, fixed in 5% neutral buffered formalin. Flow cytometric data were acquired within one week of staining using a MACSQuant 10 (Miltenyi Biotech) flow cytometer and data were analyzed using Flowjo software V10.2 (Treestar Inc.).

#### Gene expression array data mining and Ingenuity IPA analysis

Publicly available gene expression datasets (GSE24206) were mined from NCBI’s geo datasets database. Groups were defined as follows – IPF lung explants vs normal lung and IPF lung biopsies versus normal lungs. Gene expression values were extracted using NCBI’s Geo2R gene expression analysis tool and the expression data were uploaded onto ingenuity IPA. Ingenuity IPA was set to only consider changes in gene expression of 1.5-fold or greater and p ≤ 0.05. The resulting data were then overlaid on a custom designed T cell marker and immune checkpoint receptor and ligand network.

#### Statistical analysis

All statistical analyses were performed using GraphPad Prism software (GraphPad).

#### Study approval

Institutional Review Board approval for the studies outlined herein was obtained at Cedars-Sinai Medical Center. Informed consent was obtained from all patients prior to inclusion in the studies described herein. Cedars-Sinai Medical Center Department of Comparative Medicine approved all mouse studies described herein. All studies were performed in accordance with the relevant State and Federal guidelines and regulations.

## Notes

The authors have declared that no conflict of interest exists.

## References

1. du Bois, R.M. Strategies for treating idiopathic pulmonary fibrosis. Nat Rev Drug Discov 9, 129-140 (2010).

2. Hecker, L. & Thannickal, V.J. Nonresolving fibrotic disorders: idiopathic pulmonary fibrosis as a paradigm of impaired tissue regeneration. Am J Med Sci 341, 431-434 (2011).

3. Galati, D., et al. Peripheral depletion of NK cells and imbalance of the Treg/Th17 axis in idiopathic pulmonary fibrosis patients. Cytokine 66, 119-126 (2014).

4. Herazo-Maya, J.D., et al. Peripheral blood mononuclear cell gene expression profiles predict poor outcome in idiopathic pulmonary fibrosis. Sci Transl Med 5, 205ra136 (2013).

5. Kotsianidis, I., et al. Global impairment of CD4+CD25+FOXP3+ regulatory T cells in idiopathic pulmonary fibrosis. Am J Respir Crit Care Med 179, 1121-1130 (2009).

6. Reilkoff, R.A., et al. Semaphorin 7a+ regulatory T cells are associated with progressive idiopathic pulmonary fibrosis and are implicated in transforming growth factor-beta1-induced pulmonary fibrosis. Am J Respir Crit Care Med 187, 180-188 (2013).

7. Todd, N.W., et al. Lymphocyte aggregates persist and accumulate in the lungs of patients with idiopathic pulmonary fibrosis. J Inflamm Res 6, 63-70 (2013).

8. Gilani, S.R., et al. CD28 down-regulation on circulating CD4 T-cells is associated with poor prognoses of patients with idiopathic pulmonary fibrosis. PLoS One 5, e8959 (2010).

9. Moore, B.B., et al. Inflammatory leukocyte phenotypes correlate with disease progression in idiopathic pulmonary fibrosis. Front Med 1(2014).

10. Feghali-Bostwick, C.A., et al. Cellular and humoral autoreactivity in idiopathic pulmonary fibrosis. J Immunol 179, 2592-2599 (2007).

11. Lympany, P.A., et al. T-cell receptor gene usage in patients with fibrosing alveolitis and control subjects. Eur J Clin Invest 29, 173-181 (1999).

12. Shimizudani, N., et al. Conserved CDR 3 region of T cell receptor BV gene in lymphocytes from bronchoalveolar lavage fluid of patients with idiopathic pulmonary fibrosis. Clin Exp Immunol 129, 140-149 (2002).

13. Kahloon, R.A., et al. Patients with idiopathic pulmonary fibrosis with antibodies to heat shock protein 70 have poor prognoses. Am J Respir Crit Care Med 187, 768-775 (2013).

14. Shen, D.T., Ma, J.S., Mather, J., Vukmanovic, S. & Radoja, S. Activation of primary T lymphocytes results in lysosome development and polarized granule exocytosis in CD4+ and CD8+ subsets, whereas expression of lytic molecules confers cytotoxicity to CD8+ T cells. J Leukoc Biol 80, 827-837 (2006).

15. Vallejo, A.N. CD28 extinction in human T cells: altered functions and the program of T-cell senescence. Immunol Rev 205, 158-169 (2005).

16. Berg, M. & Zavazava, N. Regulation of CD28 expression on CD8+ T cells by CTLA-4. Journal of leukocyte biology 83, 853-863 (2007).

17. Rudd, C.E., Taylor, A. & Schneider, H. CD28 and CTLA-4 coreceptor expression and signal transduction. Immunological reviews 229, 12-26 (2009).

18. Weng, N.P., Akbar, A.N. & Goronzy, J. CD28(-) T cells: their role in the age-associated decline of immune function. Trends Immunol 30, 306-312 (2009).

19. Ebener, S., et al. Toll-like receptor 4 activation attenuates profibrotic response in control lung fibroblasts but not in fibroblasts from patients with IPF. Am J Physiol Lung Cell Mol Physiol 312, L42-L55 (2017).

20. Fernandez, I.E., et al. Peripheral blood myeloid-derived suppressor cells reflect disease status in idiopathic pulmonary fibrosis. Eur Respir J 48, 1171-1183 (2016).

21. Trujillo, G., et al. TLR9 differentiates rapidly from slowly progressing forms of idiopathic pulmonary fibrosis. Sci Transl Med 2, 57ra82 (2010).

22. Hogaboam, C.M., Trujillo, G. & Martinez, F.J. Aberrant innate immune sensing leads to the rapid progression of idiopathic pulmonary fibrosis. Fibrogenesis Tissue Repair 5, S3 (2012).

23. O’Dwyer, D.N., et al. The Toll-like receptor 3 L412F polymorphism and disease progression in idiopathic pulmonary fibrosis. Am J Respir Crit Care Med 188, 1442-1450 (2013).

24. Kirillov, V., et al. Sustained activation of toll-like receptor 9 induces an invasive phenotype in lung fibroblasts: possible implications in idiopathic pulmonary fibrosis. Am J Pathol 185, 943-957 (2015).

25. Brignull, L.M., et al. Reprogramming of lysosomal gene expression by interleukin-4 and Stat6. BMC Genomics 14, 853 (2013).

26. Rincon, M. & Flavell R.A. AP-1 transcriptional activity requires both T-cell receptor-mediated and co-stimulatory signals in primary T lymphocytes. EMBO J 13, 4370-4381 (1994).

27. Strioga, M., Pasukoniene, V. & Characiejus, D. CD8+ CD28- and CD8+ CD57+ T cells and their role in health and disease. Immunology 134, 17-32 (2011).

28. Dumitriu, I.E. The life (and death) of CD4+ CD28(null) T cells in inflammatory diseases. Immunology 146, 185-193 (2015).

29. Lizée, G., et al. Harnessing the power of the immune system to target cancer. Annual review of medicine 64, 71-90 (2013).

30. Naidoo, J., Page, D.B. & Wolchok, J.D. Immune checkpoint blockade. Hematology/oncology clinics of North America 28, 585-600 (2014).

31. Pardoll, D.M. The blockade of immune checkpoints in cancer immunotherapy. Nature Reviews Cancer 12, 252-264 (2012).

32. Okazaki, T. & Honjo, T. PD-1 and PD-1 ligands: from discovery to clinical application. International immunology 19, 813-824 (2007).

33. Keir, M.E., Butte, M.J., Freeman, G.J. & Sharpe, A.H. PD-1 and its ligands in tolerance and immunity. Annual review of immunology 26, 677-704 (2008).

34. Arasanz, H., et al. PD1 signal transduction pathways in T cells. Oncotarget (2017).

35. Prado-Garcia, H., Romero-Garcia, S., Puerto-Aquino, A. & Rumbo-Nava, U. The PD-L1/PD-1 pathway promotes dysfunction, but not “exhaustion”, in tumor-responding T cells from pleural effusions in lung cancer patients. Cancer Immunology, Immunotherapy 66, 765-776 (2017).

36. Asano, T., et al. PD-1 modulates regulatory T-cell homeostasis during low-dose interleukin-2 therapy. Blood 129, 2186-2197 (2017).

37. Gotot, J., et al. Regulatory T cells use programmed death 1 ligands to directly suppress autoreactive B cells in vivo. Proc Natl Acad Sci U S A 109, 10468-10473 (2012).

38. Park, H.J., et al. PD-1 upregulated on regulatory T cells during chronic virus infection enhances the suppression of CD8+ T cell immune response via the interaction with PD-L1 expressed on CD8+ T cells. J Immunol 194, 5801-5811 (2015).

39. Schwartz, R.H. Costimulation of T lymphocytes: the role of CD28, CTLA-4, and B7/BB1 in interleukin-2 production and immunotherapy. Cell 71, 1065-1068 (1993).

40. PD-1 Blockade-Mediated Rescue of Exhausted T Cells Requires CD28. Cancer Discov 7, 453 (2017).

41. Robinson, M.D. & Smyth, G.K. Small-sample estimation of negative binomial dispersion, with applications to SAGE data. Biostatistics 9, 321-332 (2008).

42. Robinson, M.D., McCarthy, D.J. & Smyth, G.K. edgeR: a Bioconductor package for differential expression analysis of digital gene expression data. Bioinformatics 26, 139-140 (2010).

43. Huang da, W., Sherman, B.T. & Lempicki, R.A. Systematic and integrative analysis of large gene lists using DAVID bioinformatics resources. Nat Protoc 4, 44-57 (2009).

44. Huang da, W., Sherman, B.T. & Lempicki, R.A. Bioinformatics enrichment tools: paths toward the comprehensive functional analysis of large gene lists. Nucleic Acids Res 37, 1-13 (2009).

